# Long-read metagenomic sequencing reveals novel lineages and functional diversity in urban soil microbiome

**DOI:** 10.64898/2026.03.20.713087

**Authors:** Yiqian Duan, Anna Cuscó, Yaozhong Zhang, Juan S. Inda-Díaz, Chengkai Zhu, Alexandre Areias Castro, Xinrun Yang, Jiabao Yu, Gaofei Jiang, Xing-Ming Zhao, Luis Pedro Coelho

**Affiliations:** College of Biomedical Engineering, Fudan University, Shanghai, China; Institute of Science and Technology for Brain-Inspired Intelligence, Fudan University, Shanghai, China; Key Lab of Organic-based Fertilizers of China and Jiangsu Provincial Key Lab for Solid Organic Waste Utilization, Nanjing Agricultural University, Nanjing 210095, China; Department of Neurology, Zhongshan Hospital, Fudan University, Shanghai, China; State Key Laboratory of Medical Neurobiology, Institutes of Brain Science, Fudan University, Shanghai, China; MOE Key Laboratory of Computational Neuroscience and Brain-Inspired Intelligence, and MOE Frontiers Center for Brain Science, Fudan University, Shanghai, China; Centre for Microbiome Research, School of Biomedical Sciences, Queensland University of Technology, Translational Research Institute, Woolloongabba, Queensland, Australia

## Abstract

City parks and other urban green spaces can bring significant benefits to the physical and mental health of city residents. However, there is limited knowledge about the microbial communities inhabiting these urban soils. Here, we applied long-read metagenomic sequencing to 58 urban soil samples from two major cities in China, enabling genome-resolved reconstruction of microbial diversity at unprecedented contiguity. We recovered 7,949 medium- and high-quality metagenome-assembled genomes, comprising 4,171 species-level genome bins, of which over 97% represent previously undescribed species. Long-read assemblies revealed extensive secondary metabolic capacity, including more than 30,000 biosynthetic gene clusters, which were highly contiguous compared with those from fragmented short-read assemblies. Beyond secondary metabolism, we uncovered over 2 million small protein families, including hundreds that are strongly enriched in the neighbourhood of defense systems and mobile genetic elements, highlighting their overlooked role in urban soils. These findings expand our understanding of the functional diversity of urban soil microbiomes and provide new insights with implications for urban public health.

## Introduction

Urban soils include parks, roadsides, sports fields, and transportation facilities. Together, they cover only approximately 3% of the global land surface^1^, yet they play an outsized role in human health by hosting microbiomes that help prevent allergies and autoimmune diseases^2–4^. Meanwhile, human activity influences soil properties, vegetation, and associated microbial communities in a reciprocal fashion^5^. Nevertheless, most soil microbiome research has focused on natural and agricultural soil ecosystems. Our understanding of the microbial composition of urban soils to which people are exposed in their daily lives remains limited^2,6^. Given that soil microbial communities exhibit greater diversity than those of other environments^7^, and that their gene pool is still far from fully sequenced^6,8^, in-depth exploration of the urban soil microbiome could uncover enormous functional potential.

Prior metagenomic studies of urban soil have mostly used short-read sequencing, which is constrained by technical limitations such as GC content bias and often yields highly fragmented assemblies^9^. In contrast, long-read sequencing technologies generate reads of sufficient length to span complex and repetitive genomic regions and thereby improve the completeness of metagenome-assembled genomes (MAGs)^10^. The availability of near-complete genomes enables the identification of complex components, such as complete biosynthetic gene clusters (BGCs)^11,12^, which are typically fragmented in short-read assemblies, as well as mobile genetic elements, including circular plasmids^13^, which are essential for microbial adaptation through horizontal gene transfer.

Here, we used long-read sequencing (Oxford Nanopore Technologies) to investigate the composition and functional diversity of microbial communities in urban soils from university campuses and parks located in two major cities (Shanghai and Nanjing) in China. Our findings revealed extensive phylogenetic diversity and novel microbial lineages, substantially expanding the known soil microbial tree of life. In addition, we systematically characterized the secondary metabolic potential of the communities, identifying over 30,000 BGCs with significantly higher contiguity than is possible with short-read based assemblies. Furthermore, we identified a diverse repertoire of small proteins, particularly those associated with defense systems and mobile genetic elements, revealing their potential roles in microbial adaptation. Overall, we provide a comprehensive resource and new insights into the microbial ecology of urban soils and its implications for public health.

## Results

### Different microbial compositions of soil samples between cities

We obtained a total of 3.28 Tbp metagenomic data via long-read sequencing (Oxford Nanopore Technology) and 3.35 Tbp via short-read sequencing (Illumina) from soil samples collected from urban green spaces, including university campuses and public parks, across two major cities (Shanghai and Nanjing) in eastern China (Fig. 1a, Supplementary Fig. 1a-c).

**Fig. 1.**
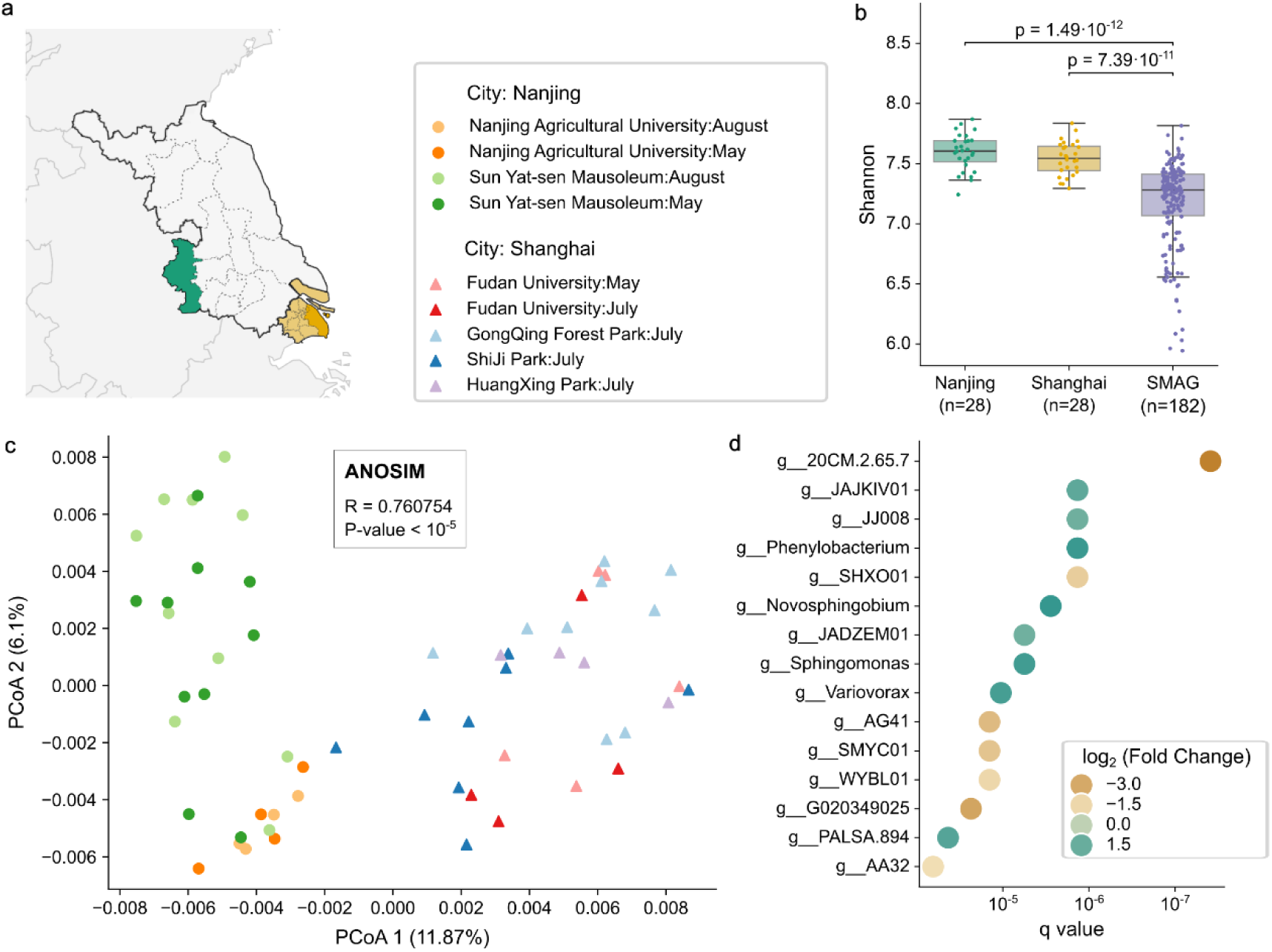
Microbial community composition and diversity in urban soils. **(a)** Graphical representation of samples from two major cities in eastern China, Nanjing and Shanghai, where samples were collected. **(b)** Comparison of the Shannon index between the two cities and samples from a publicly available soil catalog^14^. *P*-values shown are from Mann-Whitney test (two-sided alternative hypothesis), there is no statistical difference between samples from the two cities (P>0.05). Box plots indicate median (middle line), 25th, 75th percentile (box), and 5th and 95th percentile (whiskers). **(c)** Principal Coordinates Analysis of the beta diversity using the Bray-Curtis distance matrix shows differences between cities. *P*-value shown is from the ANOSIM Test (>99,999 permutations). **(d)** Significantly different abundant genera between cities identified by Maaslin3 (q-value < 0.1).

The bacterial diversity of urban soil microbiomes is comparable between the two cities studied, but was slightly higher than that of soil samples represented in the soil genome catalog (SMAG)^14^ (Fig. 1b; Supplementary Fig. 1d–e). Within the specific timeframe of this study, these soil communities exhibited significant short-term stability; samples collected from the same location at three-month intervals showed highly similar microbial compositions (Supplementary Fig. 2). Beyond this temporal consistency, the communities were structured primarily by geographic region, forming two distinct clusters for Shanghai and Nanjing (ANOSIM test, *R* = 0.76, *p*-value < 10^−5^, based on Bray-Curtis distance, Fig. 1c, Supplementary Fig. 3a-b). When considered individually, 280 microbial genera were identified as differentially abundant between the two cities (FDR < 0.1; Fig. 1d). Soil phosphorus content differed significantly between Nanjing and Shanghai, whereas the other physicochemical properties showed no notable variation (Supplementary Fig. 3c, Supplementary Fig. 4). Taken together, these results demonstrate that geographic location, coupled with local environmental factors such as the soil phosphorus content, collectively shape the taxonomic and functional profile of the urban soil microbiome.

### Construction of high-quality MAGs of urban soil

To comprehensively explore the microbial diversity of the urban soil microbial community, we used metaFlye^15^ to perform metagenomic assembly from long reads, using short reads for polishing (Fig. 2a and Methods). This resulted in 5.01 million contigs with an average N50 of 74,258 bp (Supplementary Fig. 5). Using SemiBin2^16^, we constructed 101,540 metagenome-assembled genomes (MAGs).

**Fig. 2.**
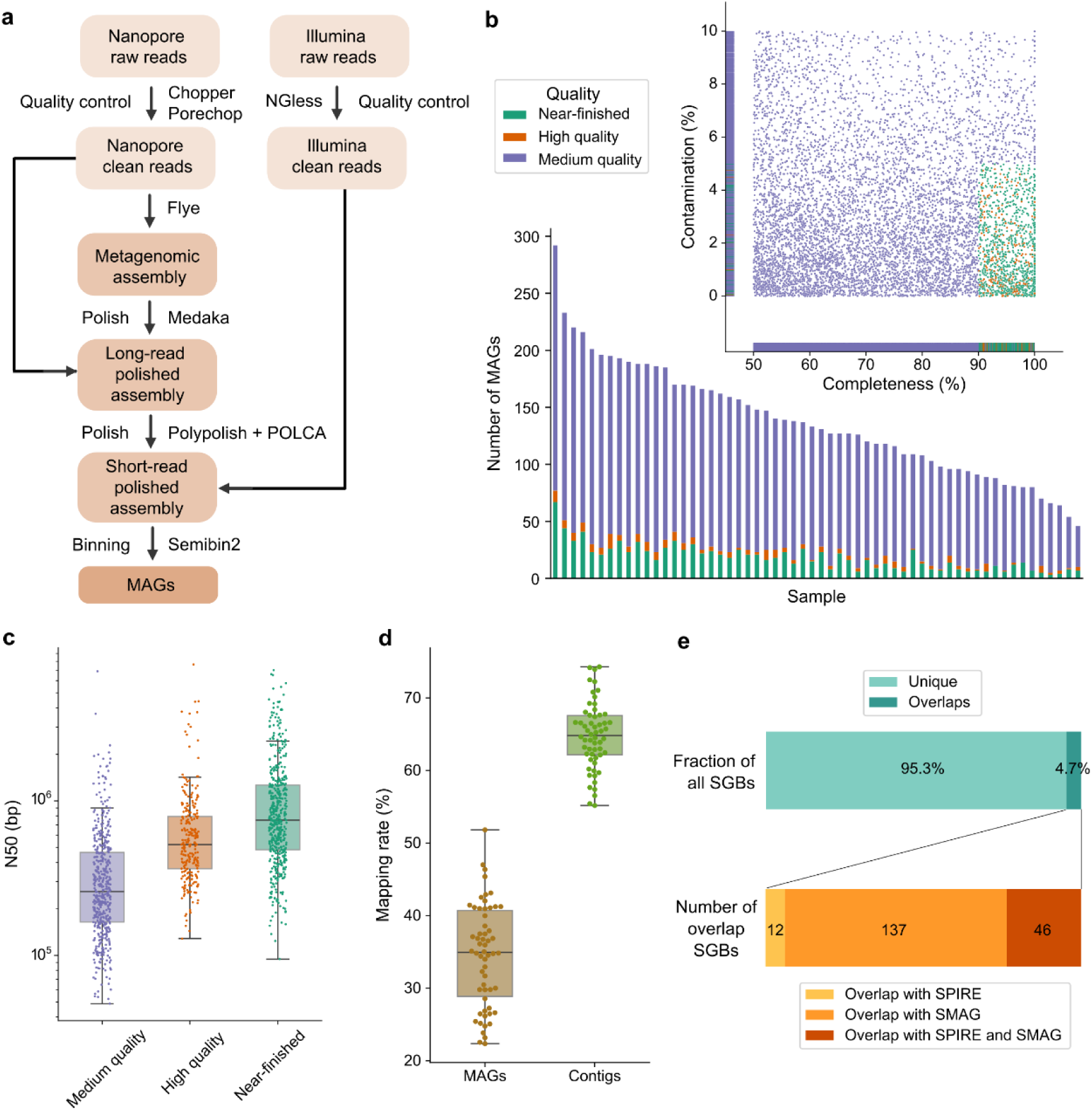
Construction and quality assessment of 7,949 metagenome-assembled genomes(MAGs) from urban soil metagenomes. **(a)** Workflow of assembly and binning using long and short reads (Methods). **(b)** The distribution of 7,949 medium-quality to near-finished MAGs among samples. A maximum of 78 high-quality or near-finished MAGs can be obtained from a single sample. The quality assessment (i.e., completeness and contamination statistics) of the 7,949 MAGs. Most medium-quality MAGs have a contamination rate of less than 5%. Each point represents one MAG, colored by class: *medium quality MAGs*: completeness > 50%, contamination < 10%, and not flagged as chimeras; Red *high quality MAGs*: completeness > 90%, contamination < 5%, and not flagged as chimeras; *near-finished MAGs*: high-quality MAGs containing at least a 5S, 16S, and 23S rRNA and at least 18 standard tRNAs, thus meeting the MIMAG standard. **(c)** Comparison of the N50 lengths of medium-, high-quality, and near-finished MAGs. For visualization, only a maximum of 500 dots per group are shown. **(d)** Mapping rate of short reads against 7,949 MAGs and all contigs. Box plots indicate median (middle line), 25th, 75th percentile (box), and 5th and 95th percentile (whiskers). **(e)** Comparison of 4,171 species-level genome bins (SGBs) against MAGs from the soil environment of the SPIRE database and the SMAG catalog.

Among all the MAGs, 6,635 MAGs were considered as medium-quality (MQ) MAGs with completeness > 50% and contamination < 10%, and 1,314 MAGs were considered as high-quality (HQ) MAGs with completeness > 90% and contamination < 5% (Fig. 2b). Among these high-quality MAGs, a total of 1,060 MAGs (80%) met the high-quality criteria as defined by the Minimal Information about a Metagenome-Assembled Genome (MIMAG) standards^17^, containing 5S, 16S, and 23S rRNA genes and at least 18 standard tRNAs, which were considered as near-finished MAGs. In total, these 7,949 medium-quality and high-quality MAGs had an average size of 4.45 Mbp (0.38 Mbp to 15.64 Mbp), an average GC content of 63.8% (28.2% to 75.6%), and an average N50 of 0.48 Mbp (23.4 kbp to 12.1 Mbp) (Fig. 2c).

To assess the representation of our MAGs within the whole metagenomes from urban soil samples, we mapped the short reads to these 7,949 MAGs and to all contigs (noting that the short reads had been used to polish, but not during the assembly process). On average, 34.52% (range from 22.37% to 51.83%) of the reads could be mapped to the MAGs. The average mapping rate increased to 64.77% (range from 55.18% to 74.29%) when mapping against all contigs including unbinned ones (Fig. 2d). This demonstrates that the MAGs represent a substantial, recoverable portion of the microbial community, while the remaining unmapped reads indicate the presence of unexplored microbial diversity^18^.

All MAGs were dereplicated and clustered into 4,171 species-level genome bins (SGBs) using dRep^19^ at 95% average nucleotide identity (ANI). Furthermore, we compared these SGBs with SMAG catalog^6^ and MAGs from the soil environment in the SPIRE database^20^. The vast majority (95.3%) of our SGBs are unique (Fig. 2e). Moreover, we used fetchMGs^21^ to extract marker genes from these MAGs and aligned them by BLASTN^22^. The results were consistent in that only a few hundred marker genes in our SGBs matched those in other reference MAGs at 95% nucleotide identity and 90% coverage (Supplementary Fig. 6a).

To further quantify the value of the long-read sequencing, we built MAGs using only the short reads from the same samples, using MEGAHIT^23^ and SemiBin2^16^ (Methods). Compared with these MAGs, the MAGs derived from long reads presented higher N50 values; fewer contigs per MAG; more unique tRNAs; and higher proportions of 16S, 23S, and 5S rRNA genes (Supplementary Fig. 6b-e).

### Species-level genome bins (SGBs) reveal phylogenetic diversity and novel lineage references

To further investigate the phylogenetic diversity of the obtained SGBs, we used GTDB-tk^24^ to assign taxonomy to the 4,171 SGBs. Most bacterial species (97.7%) are novel, since only 138 (3.3%) of the SGBs could be assigned to previously described species, and 17.5% of the SGBs represent novel genera (Fig. 3a). The vast majority of SGBs were classified as bacteria, with archaea representing only a minor fraction (0.94%). The dominant phyla were Acidobacteriota (*n* = 1,251) and Pseudomonadota (*n* = 950), followed by Gemmatimonadota (*n* = 282)(Fig. 3b-c).

**Fig. 3.**
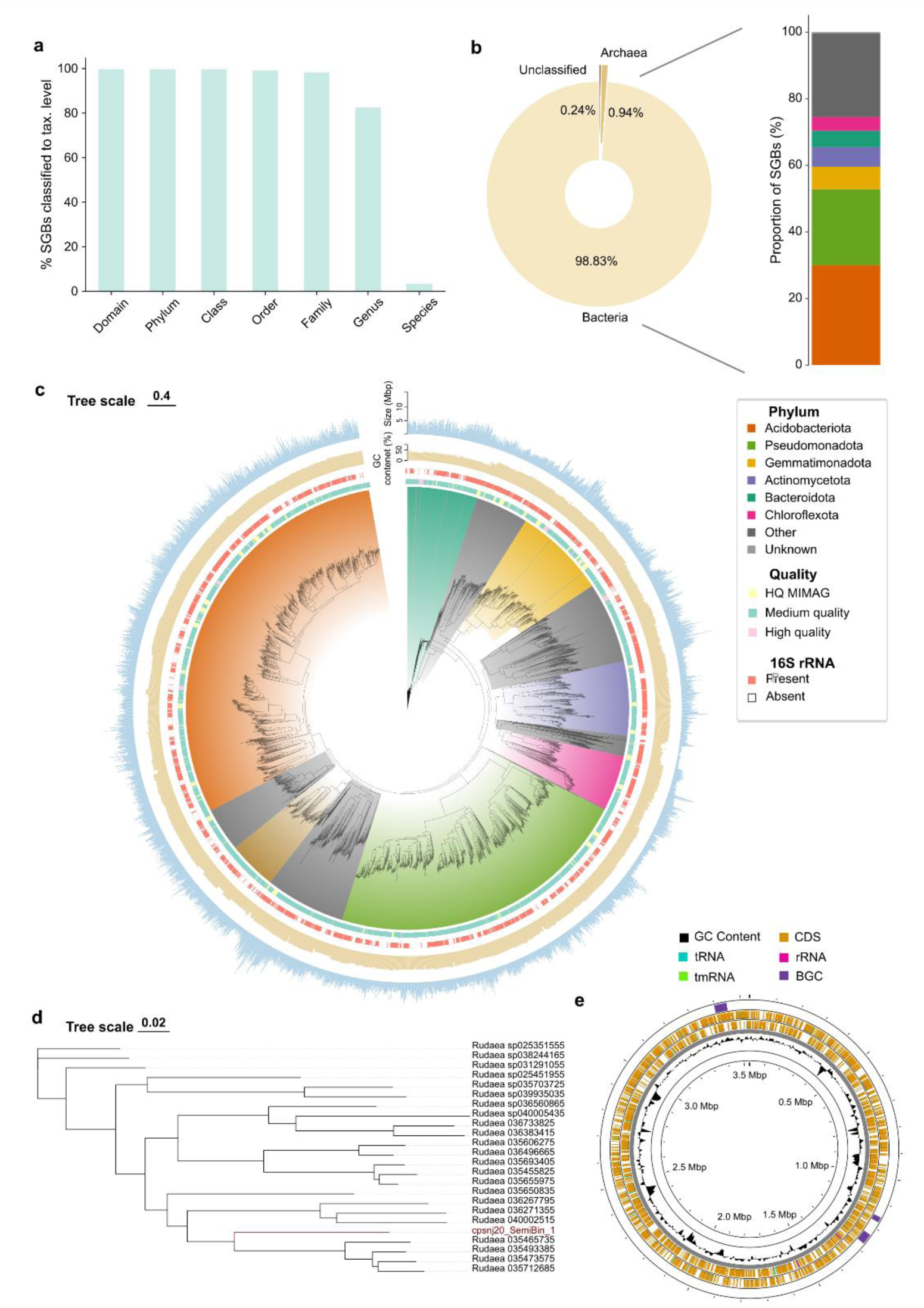
Phylogenetic diversity and novel species discovery in urban soil microbiome. **(a)** Taxonomic assignment of the 4,171 SGBs from urban soil based on the GTDB database. **(b)** Taxonomic classification rates of SGBs at different taxonomic levels. **(c)** Phylogenetic tree of the 4,171 SGBs showing size, GC content, quality assessment, and the presence of 16S rRNA sequences. **(d)** Phylogenetic tree of the longest circular MAG and representative genomes from species of the genus *Rudaea*. **(e)** The longest circular MAG (3.53 Mb) is shown, annotated with the GC content, CDS, tRNA, rRNA, tmRNA, and BGC regions.

A total of 648 SGBs were represented by high-quality MAGs; among these, 601 were not assigned to species-level annotation by GTDB-tk^24^. Thus, we attempted to extract complete 16S rRNA genes from these SGBs and match them to the MicrobeAtlas database of operational-taxonomic units (OTUs)^25^ using MAPseq^26^ (Methods). We found that 232 out of the 601 SGBs (38.6%) did not have close matches (≥ 97% identity) to any sequences in the MicrobeAtlas database^25^, indicating taxa that were not previously reported. In addition, 179 SGBs (29.8%) contained 16S rRNA genes with 99% identity matches to OTUs in the MicrobeAtlas database^25^, but these OTUs either lacked reference genomes or could not be linked to genomes in the Progenomes2 database^27^. Our MAGs can, therefore, provide genomic context for these underrepresented OTUs, directly associating specific functions with OTUs.

Among the 47 SGBs that did receive a species-level annotation from GTDB-Tk, 24 SGBs contain complete 16S rRNA genes while their representatives in the GTDB database r226 lack them^28^. These results indicate that long-read sequencing enables the assembly of more contiguous genomes, thereby facilitating the recovery of full-length 16S rRNA genes. This allows for supplementing existing databases, providing associations between functionality and OTUs, and revealing undescribed microbial diversity.

In total, we obtained 55 SGBs whose representative MAG consists of only a single contig, 38 of which were near-finished MAGs that met MIMAG standards^17^. Among these 38 MAGs, 35 lack species-level annotations. Notably, one of these 35 unannotated MAGs, CSMAG_0101, was the longest (3.53 Mb) circular MAG with 99.99% completeness, and was assigned to the genus *Rudaea*. Comparative genomic analysis against all the reference genomes from *Rudaea* species in the GTDB database^28^ indicates that this SGB constitutes a novel species within the genus as evidenced by an ANI < 95% and phylogenetic reconstruction (Fig. 3d). *Rudaea* species are known for their ability to decompose cellulose, which may have potential applications in biomass utilization, such as straw degradation and composting treatment^29^. Three biosynthetic gene clusters (BGCs) for putative arylpolyene, hserlactone, and hybrid NRPS/T1PKS compounds were identified from this MAG (Fig. 3e), revealing a biosynthetic capacity of this novel species of *Rudaea*, that may facilitate the environmental adaptation for microbial competition in urban soil.

The remaining three SGBs with species annotation, probably representing novel strains based on their distinction from their closest genomic relatives in the GTDB database^28^: DASTBI01 sp035712145 (ANI = 95.9%), *Nitrospira C* sp029240595 (ANI = 96.5%), and *Rhodanobacter* sp035571995 (ANI = 98.7%). The representative MAG for these three SGBs also exhibited improvements in genome completeness and/or a reduction in contamination relative to the corresponding reference genomes: DASTBI01 sp035712145 (completeness: 99.37% vs. 98.74%; contamination: 0.07% vs. 1.14%), *Nitrospira C* sp029240595 (completeness: 99.82% vs. 85.49%; contamination: 0.49% vs. 0.42%), and *Rhodanobacter* sp035571995 (completeness: 91.39% vs. 90.06%; contamination: 0.11% vs. 1.53%).

### Discovery of extensive secondary metabolite biosynthetic potential from MAGs

Secondary metabolites have been identified in previous metagenomic studies by short-read sequencing^6,18,30^, however, the fragmented nature of short-read assemblies often fails to capture full-length sequences. Long-read sequencing enables a better description of the biosynthetic potential in complex microbiomes by facilitating the recovery of contiguous, high-quality genomic regions^11,12,31,32^.

Here, we recovered 31,634 BGCs, which clustered into 16,301 gene cluster families (GCFs). These BGCs were identified in approximately 94% (7,458 out of 7,949) of the medium- and high-quality MAGs assembled from long reads. Following the classification criteria of Genomes from Earth’s Microbiomes (GEM) catalog^18^ and the SMAG catalog^6^, we categorized the identified BGCs by their genomic features.

Specifically, we identified 11,340 (35.8%) large and modular BGC regions exceeding 40 Kb and 18,082 (57.2%) complex BGCs (10-30 Kb) harboring multiple genes (Fig. 4a). Compared with the BGCs derived from short-read–assembled MAGs in the SMAG catalog^6^, our long-read-assembled MAGs contained a significantly lower proportion of fragmented BGCs (< 10 Kb) (0.45% vs. 35.9%; *p*-value < 10^−308^, chi-square test).

**Fig. 4.**
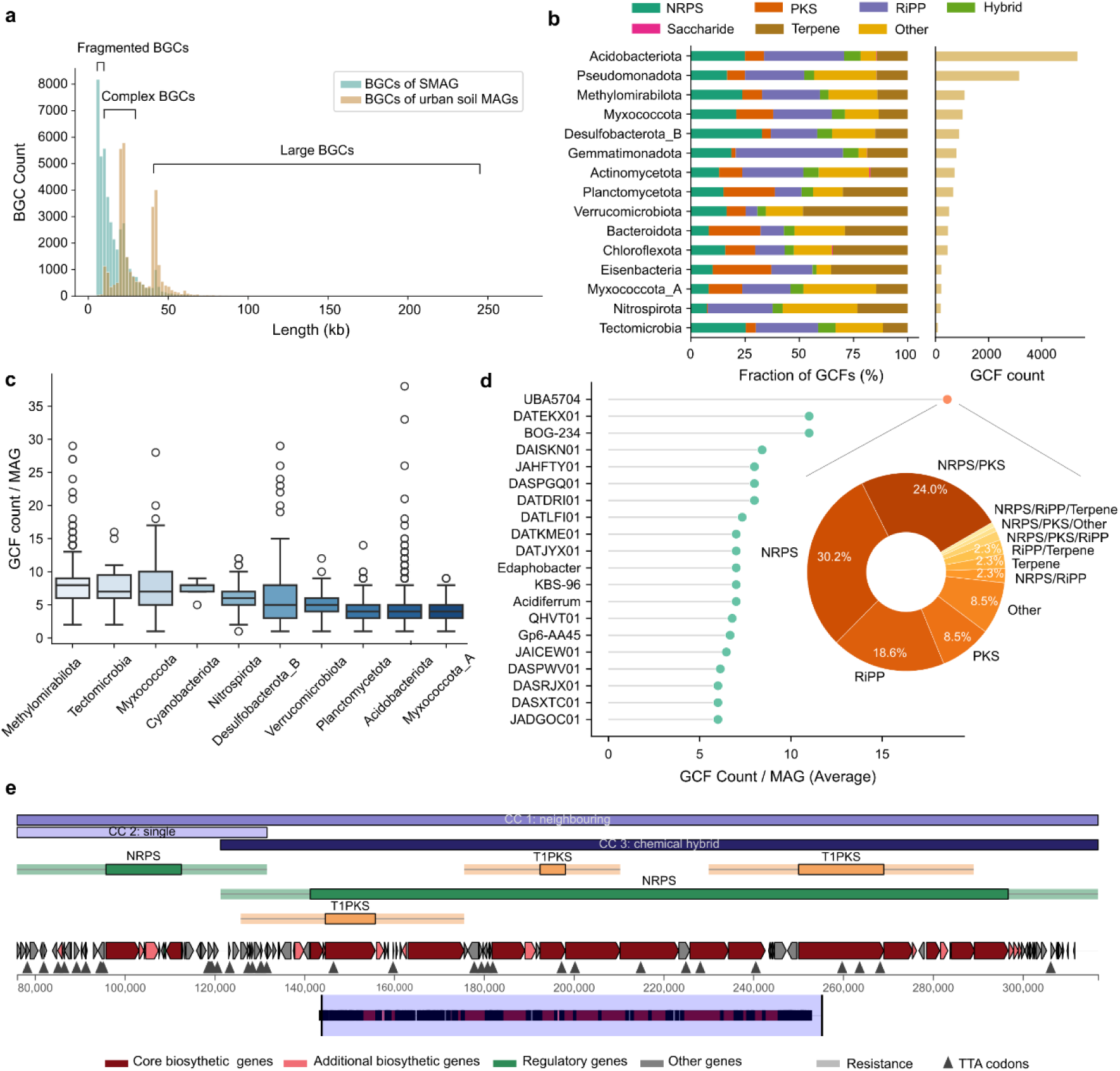
Biosynthetic gene cluster diversity in SGBs. **(a)** Comparison of BGC length between BGCs of urban soil MAGs and those of the SMAG catalog. **(b)** The top 15 phyla with the most biosynthetic gene cluster families (GCFs), and the composition of the GCFs in each phylum. NRPS: Non-ribosomal peptide synthetase; RiPP: Ribosomally synthesized and post-translationally modified peptide; PKS: polyketide synthase. **(c)** The top 10 phyla with the highest GCF count per MAG. Box plots indicate median (middle line), 25th, 75th percentile (box), and 5th and 95th percentile (whiskers) as well as outliers (single points) that lie beyond 1.5 IQRs of the lower and upper quartile. **(d)** The top 20 genera of Acidobacteriota with the highest density of GCFs and the distribution of GCF categories in the genus UBA5704. **(e)** The composition of the largest BGC region from the genus UBA5704.

Among these 16,301 GCFs, ribosomally synthesized and post-translationally modified peptides (RIPPs) constituted the most abundant category (n = 4760, 29.2%) and were distributed across 37 phyla (Fig. 4b). The vast majority (97.4%) of the GCFs identified in Thermoproteota (Archaea) were classified as RIPP gene clusters. Nonribosomal peptide synthetases (NRPS) gene clusters (20.1%) and terpene gene clusters (17.8%) were also widely represented, originating from 26 and 29 phyla, respectively.

Within the whole microbial community, Acidobacteriota, Pseudomonadota, and Methylomirabilota collectively contributed more than 58% of all GCFs (Fig. 4b). Further, we assessed the biosynthetic production capacity of each phylum. On average, each phylum contained 4.22 ± 2.97 GCFs per MAG (Fig. 4c). The highest secondary metabolic capability was observed in Methylomirabilota (8.19 BGCs per MAG), Tectomicrobia (7.80 BGCs per MAG), and Myxococcota (7.66 BGCs per MAG). Recent genomic studies suggest that large genomes and predatory lifestyles drive the expansion of BGCs in Myxococcota^33^. The high biosynthetic capacity observed in Tectomicrobia^34^ and Methylomirabilota^35^ further highlights the importance of discovering novel bioactive molecules from uncultured “microbial dark matter” that may mediate unique ecological processes.

Although Acidobacteriota possesses the most GCFs, the biosynthetic potential of its various genera is unevenly distributed (Fig. 4d). Consistent with previous reports^6,36^, UBA5704 shows extremely high biosynthetic potential (18.57 GCFs per MAG), whereas 69 genera (55.2%) have a biosynthetic potential lower than the average for all phyla (4.22). Among the 128 GCFs of UBA5704, the NRPS (30.2%) and hybrid NRPS/PKS (24.0%) gene clusters were predominant. In particular, CSMAG_5610 exhibited 38 BGC regions, including 14 NRPS and 12 hybrid NRPS/PKS gene clusters. Notably, we identified the longest and complete BGC region (240,746 bp) from UBA5704, encoding 16 biosynthetic genes and 15 biosynthetic-additional genes, containing 36 NRPS/PKS modules (Fig. 4e). Its protocluster is similar to BGC0002135.2 in the MIBiG database^37^, which is from *Xenorhabdus bovienii* SS-2004 and produces bovienimide A.

### Defense system related small proteins

Although small proteins, under 100 amino acids in length, have historically been overlooked^38^, they are increasingly recognized as crucial players in the microbial community^39,40^. To investigate these molecules, we used the GMSC-mapper^40^ to predict small open reading frames (smORFs) homologous to the Global Microbial smORFs Catalog (GMSC)^40^ from the 7,949 medium-quality and high-quality MAGs and unbinned contigs. A total of 6,356,825 smORFs were predicted, resulting in 4,497,401 non-redundant smORFs after dereplication (Supplementary Fig. 7a).

Among these smORFs, 35.72% could be annotated at the genus or species level (Supplementary Fig. 7b). These smORFs were subsequently clustered at 90% amino acid identity and 90% coverage using CD-HIT^41^, yielding 2,206,901 small protein families (Supplementary Fig. 7c). Furthermore, conserved domain annotation was performed using RPS-BLAST^42^ against the Conserved Domain Database (CDD)^43^, which revealed that only 354,271 families (16.05%) contained known domains (Supplementary Fig. 7d), indicating that the vast majority of small protein families remain functionally uncharacterized.

Therefore, we focused on the 1,852,630 small protein families lacking conserved domain annotations. Given previous evidence that persistent interactions between phages and hosts in soil drive the evolution of diverse bacterial defense mechanisms, we screened for small proteins potentially associated with defense systems^44,45^. Using a curated COG (Clusters of Orthologous Groups)/Pfam (Protein Families) list of 427 known defense system related genes from a previous study^46^, we identified small protein families located within 10 genes upstream or downstream of these defense genes. A total of 99,389 (5.36%) unannotated small protein families were identified as potentially defense-related, with at least one member located near a known defense gene. We further investigated 9,418 small protein families containing 8 or more members; among them, 656 (6.96%) had at least half of their members associated with defense systems. Notably, within Family_1682595, 30 out of 31 members (96.8%) were defense-related. This represents a significant enrichment compared with a genomic background frequency of 4.45% for defense-related genes (*p*-value = 8.63 · 10^−40^, binomial exact test). This family encodes a small protein of 51 amino acids, distributed across multiple species of *Nitrospira*^47^, and has a conserved genomic context and sequence conservation (Fig. 5a-b).

**Fig. 5.**
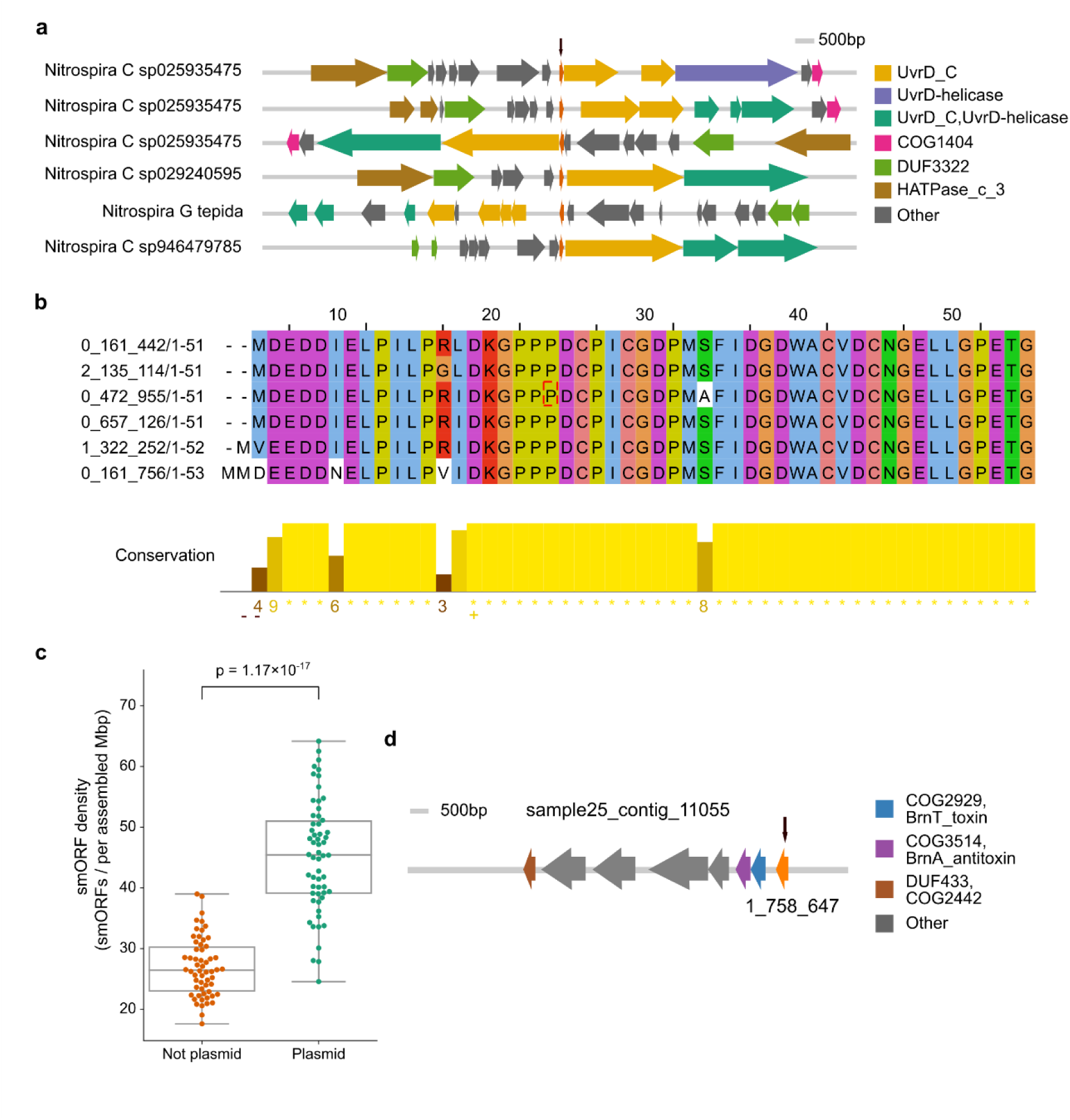
Defense system-related small proteins. **(a)** A family of small proteins (Family_1682595) that are potentially involved in the UVR system in the stress defense of *Nitrospira C*. Thirty out of 31 members of this family exhibit conserved gene neighborhoods, located downstream of two-component signal transduction system-related genes (HATPase_c_3) and directly upstream of UVR system-related genes (UvrD_c, UvrD-helicase). Only the genomic neighborhoods of the 6 members that have been annotated to the species level are displayed. **(b)** High sequence conservation in small protein family Family_1682595 as shown by multiple sequence alignment. **(c)** The density of small proteins found within plasmids and those not found in plasmids was compared in each sample. Box plots indicate median (middle line), 25th, 75th percentile (box), and 5th and 95th percentile (whiskers). *P*-values shown are from the Mann-Whitney test (two-sided). **(d)** The small protein (1_758_647) in a circular plasmid is directly upstream of the BrnT/BrnA toxin-antitoxin (TA) system (BrnT_toxin and BrnA_antitoxin domains).

Specifically, it is located upstream of defense genes containing UvrD-C and UvrD helicase domains^48,49^ and downstream of defense genes with DUF3322 and HATPase_c3^50^ domains. Therefore, we hypothesize that this small protein family may serve as a signal regulator, linking the environmental signal transduction mediated by HATPase_c3 and participating in DNA repair mediated by UvrD. It could potentially enhance the stress defense and genomic stability of *Nitrospira* and maintain nitrification in diverse environments, such as wastewater, soil, and freshwater habitats.

Long-read sequencing also uniquely enables the complete reconstruction of circular elements^13^. We obtained 11,679 circular contigs, of which only 130 (1.11%) were considered full MAGs and exhibited a longer average length than the remaining circular contigs (Supplementary Fig. 5b-d). We also predicted 13,695 contigs as putative plasmids using geNomad^51^, including 71 circular plasmids.

Small proteins were significantly more abundant on plasmid contigs than on non-plasmid contigs (*p*-value = 1.17 · 10^−17^, Fig. 5c). Similarly, small proteins have been shown to be more prevalent in phages than in host prokaryotic genomes^52^. Together, these results indicate that the enrichment of small proteins is a common feature shared by mobile genetic elements such as plasmids and phages. Among the 71 circular plasmids, we found a small protein (CS.small.protein.100AA.000_027_965, containing 75 amino acids) with conserved genomic neighborhood patterns across multiple contigs. It is located directly upstream of a toxin-antitoxin (TA) system with COG2929 and BrnT_toxin domains, as well as a TA system with COG3514 and BrnA-antitoxin domains (Fig. 5d). The alignment result showed that this small protein shares 93% sequence identity with proteins from the DUF6429 family in Burkholderia. However, the function of this protein is still unknown. Based on its conserved genomic context, we speculate that it may constitute a component of the BrnT/BrnA toxin-antitoxin system^53^, potentially maintaining stable inheritance of plasmids.

### Characterization of the urban soil antibiotic resistome

To investigate the presence and spread of antibiotic resistance genes (ARGs) in urban soil^54^, we used the tool fARGene^55^, which is designed to predict ARGs with high sensitivity. We complemented this approach by filtering the predicted ARGs by matching them against the ResFinder database^56^ using stringent thresholds (minimum 80% nucleotide identity and 60% coverage) to identify known ARGs with high confidence. We considered the outputs from fARGene that did not meet these further stringent criteria as latent ARGs (as they may represent novel ARGs but require further evidence to confirm their function as ARGs^57^), while those that met the criteria were considered established ARGs.

We observed a very large number of latent ARGs, with a total of 15,952 ARGs predicted by fARGene (Fig. 6a, Supplementary Fig. 8). However, only 12 genes could be considered established. The most common ARG gene classes were beta-lactamases (66%) and aminoglycosides (32.3%) (Supplementary Fig. 8). In particular, the metallo-beta-lactamases (MBLs) B3 made up 53.2% of the total ARGs, consistent with previous findings that MBLs are commonly found in soil^58,59^. Most (78.2%) ARGs were found in a single sample and only 11.7% ARGs were found in samples from both cities (Fig. 6b) with the resistome composition clustering by city (Fig. 6c).

**Fig. 6.**
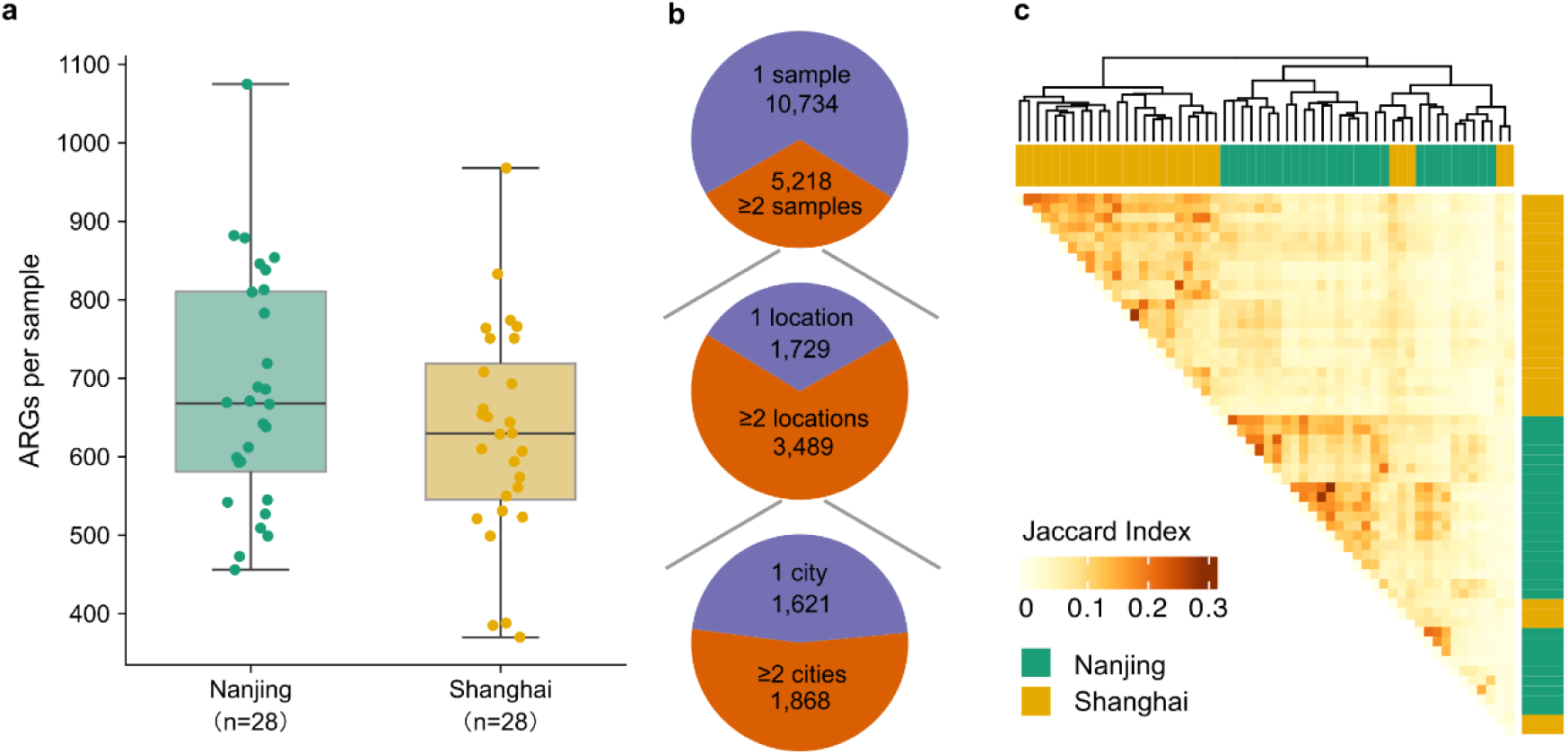
Antibiotic resistance genes. **(a)** The number of ARGs found per sample from all contigs in Nanjing and Shanghai and the distribution in both cities. Box plots indicate median (middle line), 25th, 75th percentile (box), and 5th and 95th percentile (whiskers). **(b)** Distribution of ARGs across a geographic hierarchy. Most ARGs are sample-specific; those shared across samples are stratified by co-occurrence within the same location, across locations in the same city, or across cities. Sampling times were not taken into account. **(c)** The pair-wise Jaccard’s index based on the number of ARGs found in single samples and shared between samples clusters the metagenomes by city. Hierarchical clustering was performed on the Jaccard similarity matrix using Euclidean distance and the complete linkage method.

## Discussion

Here, we constructed a comprehensive urban soil microbiome resource through long-read sequencing of soil samples collected from university campuses and parks in Shanghai and Nanjing, China. Our analyses revealed that these urban green spaces harbor extraordinary phylogenetic novelty, extensive chemical biosynthetic capacity, and a dense repertoire of small proteins associated with defense and mobility. Our findings challenge the perception of urban soils as biologically simplified systems and instead position them as dynamic reservoirs of microbial diversity.

Recent studies have indicated that, for bacterial genomes, given the extremely low error rate of nanopore R10.4.1 sequencing, the use of short reads is no longer required to obtain low error rates^60,61^. In contrast, we found that polishing with long and short reads still improved the completeness of MAGs from these complex soil samples. This is also consistent with our previous findings on the gut microbiome of pet dogs, which is a host-associated habitat^62^. These findings suggest that polishing may still benefit genome reconstruction from metagenomes of complex microbial communities.

Notably, we discovered the longest complete BGC region from the acidobacterial genus UBA5704, which has been independently reported to harbor an unusually high number of biosynthetic enzymatic domains with a much longer length^6,36^.

Subsequent exploration of the phylogenetic distribution across various habitats and the genome structure of this biosynthetic potential in the UBA5704 clade could reveal the evolution and diversification of BGCs in this bacterial lineage with unclear characteristics. These findings may also provide broader insights into the specific role of Acidobacteria in soil metabolic networks and microbial interactions.

We found a large repertoire of latent antibiotic resistance genes (ARGs) and only a small number of established ARGs in urban soils. This leads to a nuanced picture of the risks posed by these genes. On the one hand, many of these computational predictions may not be phenotypically resistant. On the other hand, the presence of a large number of latent ARGs suggests that urban soils can serve as reservoirs of resistance genes that may be mobilized in the future, and past work has shown that genes at low identity from known ARGs can confer clinically relevant resistance^58^.

We predicted small open reading frames (smORFs) from both MAGs and unbinned contigs using our previously developed GMSC-mapper^40^, which performs a homology search against the Global Microbiome smORF Catalog (GMSC)^40^. The presence of homologous sequences in GMSC, especially those found in multiple independent samples or different habitats, enhances the confidence of the predicted smORF, indicating that it is unlikely to be random genomic noise or a prediction error. Here, through gene neighbourhood analysis, we focused on characterizing novel functions of these small proteins encoded by smORFs that are homologous to GMSC entries. However, the GMSC remains far from saturated across many habitats, especially in soil, where a substantial proportion of novel smORFs remains unknown. This finding highlights the extensive functional diversity of smORFs, which encode small proteins in urban soils that have yet to be explored.

Overall, our study shows that urban soils are hotspots of microbial novelty and functional complexity. By resolving genomes, metabolites, and adaptive elements at unprecedented resolution with long-read metagenomics, we provide a foundational resource for understanding microbial ecology at the human–environment interface. As urbanization continues globally, it will be indispensable to uncover how microbial communities evolve, compete, and interact within the cities we inhabit.

## Methods

### Sample collection

A total of 30 surface soil samples (approximately 10 cm depth) were collected from the following locations in Shanghai, China: Handan campus of Fudan University, Gongqing Forest Park, Huangxing Park, and Century Park. Additionally, 28 surface soil samples were collected from the following locations in Nanjing, China: Weigang campus of Nanjing Agricultural University and Zhongshan Mausoleum National Park. Surface soil was randomly sampled within each location to ensure spatial representativeness; except in the Zhongshan Mausoleum region, samples were collected along Bo’ai Road, progressing towards the interior of the scenic area, and returning via Youju West Road. During the sampling period, the average annual temperatures in Shanghai and Nanjing are approximately 18.5°C and 17.1°C, respectively. The average annual precipitation in Shanghai and Nanjing is 1414.2 mm and 1276.8 mm, respectively. All the samples were immediately frozen in dry ice after collection and stored at −80 °C until DNA extraction.

Two pairs of samples (samples 3 & 4, and samples 11 & 12) were collected as technical replicates. In all downstream statistical analyses, only one sample from each replicated pair was retained.

### Determination of soil physicochemical properties

The soil physicochemical properties were analysed by Shanghai BIOZERON Biotechnology Co., Ltd.(Shanghai, China). The measured parameters included total carbon (TC, g/kg), soil organic carbon (SOC, g/kg), total nitrogen (TN, g/kg), total phosphorus (TP, g/kg), total potassium (TK, g/kg), and electrical conductivity (EC, μs/cm). These properties were determined following the Chinese standards for soil property analysis (available at www.chinesestandard.net): EC (HJ 802–2016), TC (HJ 695-2014), SOC (NY/T 1121.6-2006), TN (HJ 717-2014), TP (HJ 632-2011), and TK (NY/T 87-1988).

### Metagenomic sequencing

Urban soil samples were processed and sequenced by Novogene (Beijing, China). High-quality genomic DNA was extracted by the CTAB method, DNA degradation and pollution were monitored on 1% agarose gel, and the OD 260/280 ratio was detected by Nanodrop to check the purity of DNA. DNA concentration was measured using the Qubit®DNA detection kit in the Qubit®3.0 Flurometer (Invitrogen, USA).

The total amount of DNA in each sample was at least 0.2 μg, which was used as input material for DNA library preparation. According to the manufacturer’s instructions, NEB Next®Ultra™Illumina DNA Library Preparation Kit (NEB, USA) was used to generate sequencing libraries. The DNA library was sequenced on an Illumina NovaSeq 6000, resulting in 150bp pair-end reads.

According to the standard procedure provided by Oxford Nanopore Technologies (ONT), the experimental procedure mainly includes the following steps: based on the genomic DNA detection results, using Covaris g-TUBE for interruption processing, selecting DNA fragments, and preparing the DNA library using standard ligation methods. The library inspection is qualified, and sequencing will be performed using the Flow Cell (R10.4.1 chip) dedicated to the PromethION P48 instrument based on the effective concentration of the library and data output requirements (40G). Basecalling was performed on the binary fast5 format raw data containing all raw sequencing signals from Nanopore sequencing using Guppy v6.4.6 with the High-Accuracy (HAC) model, and then the fast5 format was converted to fastq format.

### Metagenomic quality control and assembly

We used Chopper v0.3.0^63^ to remove long reads with the Phred average quality score below 10, length below 500 bp, and sequences from human. Then we used Porechop v0.5.0^64^ to locate and remove the adapters. For short reads, we used NGLess v1.5^65^ to trim base pairs with quality below 25 and discard reads shorter than 45 bp. Then, we used Flye v2.9.2^15^ to assemble clean long reads with the “–nano-raw” and “–meta” parameters. We used Medaka v1.9.1 to polish assembled contigs with long reads using “r1041-e82_400bps_hac-v4.2.0” as the inference model. Subsequently, Polypolish v0.5.0^66,67^ was used to polish with clean short reads. Finally, we used the POLCA tool built into MaSuRCA v4.1.0^68,69^ to further improve the consistency quality of assemblies by utilizing short reads. For comparison, the clean short reads were also assembled with MEGAHIT v1.2.9^23^.

### Metagenomic binning and quality assessment

We used SemiBin2 v1.5.1^16^ to generate metagenome-assembled genomes (MAGs) from contigs with the soil environment selected as the pre-training model. Then we used Checkm2 v1.0.1^70^ to evaluate the completeness and contamination of MAGs.

GUNC v1.0.6^71^ was used to detect potentially chimeric MAGs. MAGs with completeness > 50%, contamination < 10%, and which passed GUNC’s chimera check were considered medium-quality MAGs. MAGs with completeness > 90%, contamination < 5%, and passing GUNC’s chimera check were considered high-quality MAGs. Subsequently, Barrnap v0.9^72^ was used to predict rRNA in MAGs. tRNAscan v2.0.12^73^ was used to search for tRNA in MAGs with “-B” for bacteria and “-A” for archaea. Among high-quality MAGs, those containing at least a 5S, 16S, and 23S rRNA and at least 18 standard tRNAs were considered to be near-finished MAGs that meet the MIMAG standard^17^. We used CoverM v0.7.0^74^ to calculate MAG coverage with “–mapper minimap2^75^ ont –methods mean”.

### Species-level genome bins (SGBs) clustering

The 7,949 medium-quality and high-quality MAGs were dereplicated and clustered into 4,171 species-level genomes (SGBs) based on 95% average nucleotide identity (ANI) by dRep v3.5.0^19^ with ‘-p 40 -comp 50 -con 10 -sa 0.95’.

### Comparison of SGBs to public databases

Our 4,171 SGBs were compared against 21,077 MAGs from the SMAG catalog^6^ and 15,975 MAGs from the soil environment in the SPIRE database^20^. All SGBs and MAGs of these databases were independently dereplicated based on 95% ANI by dRep v3.5.0^19^ with ‘-p 40 -comp 50 -con 10 -sa 0.95’ separately. Furthermore, we used fetchMGs v2.0.1^76^ to extract marker genes from all SGBs and MAGs of these databases. Based on the criteria used in mOTUs software^77^, five universal single-copy marker genes (COG0018, COG0215, COG0495, COG0525, and COG0541) were selected for comparison. These marker genes were chosen for their optimal sequence length and high alignment identity, which provide sufficient phylogenetic resolution to distinguish closely related species. Finally, these marker genes were aligned by BLAST^22^ separately; MAGs sharing at least 95% amino acid identity (AAI) across these markers were classified as the same species.

### Taxonomy assignment

We assigned taxonomy for 4,171 SGBs using GTDB-tk v2.4.1^24^ based on the GTDB r226 database^28^. We used the MMseq2 taxonomy^78,79^ to assign taxonomy to contigs that had not been binned into MAGs. The phylogenetic tree was visualized by tvBOT^80^. The bacterial genome was visualized by Proksee^81^.

### Biosynthetic gene clusters (BGCs) identification

We used antiSMASH v7.0.0^82^ to identify BGCs from MAGs. Then, we used BiG-SCAPE v2.0^83^ to group BGCs with ‘–mix -c 8 –include-singletons’.

### Gene prediction and function annotation

We used Prodigal v2.6.3^84^ to predict genes from MAGs and unbinned contigs, using “-p meta” for unbinned contigs. Eggnog-mapper^85^ was used for function annotation based on the Eggnog5.0 database^86^. E-values < 1e-5 were considered significant.

### Small protein prediction

We used GMSC-mapper v0.1.0^40^ to predict small proteins encoded by smORFs homologous to the Global Microbial smORFs Catalog (GMSC)^40^ from medium-quality and high-quality MAGs and unbinned contigs. We used CD-HIT^41^ to cluster non-redundant small proteins at 90% amino acid identity and 90% coverage, resulting in small protein families. We used RPS-BLAST^42^ to search predicted smORFs against the conserved domain database (CDD)^43^. Matches with E-values < 0.01 and coverage > 80% were considered significant.

For sequences with conserved gene neighborhoods, ClustalW^87^ was used for multiple sequence alignment, and Jalview^88^ was used to visualize the results of the multiple sequence alignment.

The density of smORFs is defined as: *ρ* = *n*_*smORFs*_ / *L*, where *n*_*smORFs*_ is the number of redundant smORFs, and *L* is the assembled length in megabase pairs (Mbp).

### Plasmid identification

We used geNomad v1.8.1^51^ to identify plasmids; contigs with plasmid prediction scores > 0.95 were considered plasmids. Among these, contigs reported as “circular” by Flye were considered circular plasmids.

If at least one member of the small protein family came from plasmids, then the small protein family was considered as being present in plasmids.

### Antimicrobial resistance genes (ARGs) identification

We identified antibiotic resistance genes (ARGs) using fARGene v0.1 with 17 hidden Markov model (HMM) profiles and default parameters within contigs. The HMMs targeted five major antibiotic classes: beta-lactams (classes A, B1/B2, B3 and D)^55,89^, aminoglycosides (*aac(2’)*, *aac(3)*, *aac(6’)*, *aph(2’‘), aph(3’)* and *aph(6)*)^90^, macrolides (*erm* and *mph*)^91^, quinolones (*qnr*)^92^, and tetracyclines (efflux pumps, monooxygenases and ribosomal protection genes)^93^. The identified open reading frames in amino acid format were clustered at 95% sequence identity using CD-HIT v4.8.1^41,94^, respectively, and downstream analyses were performed at the centroid level. Functional annotation of ARGs from unbinned contigs using eggNOG-mapper v2.1.13^85^ with the eggNOG v5.0 database^86^ showed that 83.3% of genes were annotated with antibiotic resistance functions based on Pfam, gene descriptions or best hits. A further 16.3% of genes, derived from fARGene’s *aac* models, matched acetyltransferase-associated Pfam entries. Recently described aminoglycoside families not captured in earlier models may account for these annotations^90^. To determine if a gene was established– encountered in clinical pathogens and present in current repositories –or latent^57^– computationally predicted ARGs not present in current repositories -we analyzed the open reading frames in nucleotide format with ResFinder v2.4.0^56^ with identity threshold at nucleotide level and minimum coverage at 80% and 60%, respectively. Any gene identified by ResFinder and the genes in its cluster were considered established ARGs, while the rest were considered latent ARGs. Only 12 genes in 11 clusters were considered established ARGs.

Additionally, we used Resistance Gene Identifier (RGI) v6.0.3^95^ with the – include_loose parameter against the Comprehensive Antibiotic Resistance Database (CARD 2023)^95^. RGI initially reported 40,712 putative ARGs from predicted genes on contigs, of which 95% were annotated as efflux pump *adeF* or *van* genes. As RGI predictions are alignment-based, we applied *a posteriori* filter requiring at minimum of 80% sequence identity, retaining 540 genes as putative ARGs. Among these, 274 (50.7%) matched the Mycobacterium tuberculosis *rpsL* gene. A further 185 genes (34.3%) were annotated as the efflux-associated gene *rsmA*. Due to the high proportion of low-identity hits, RGI-derived ARG predictions were excluded from downstream analyses, but the full results are available in the associated Zenodo repository.

### Alpha diversity and differential abundance analysis

Operational-taxonomic unit (OTU) tables of urban soil samples and canine gut samples^62^ were generated by SingleM^96^. OTU tables for the SMAG catalog samples (BioProject ID: PRJNA983538)^14^ were extracted from the Sandpiper^96^ database (which had been computed with the same tool). Bacterial alpha diversity indices (Chao1, Shannon, and Simpson) were computed per OTU and then aggregated at the sample level using the mean. Benchmarking between 3 different aggregation methods (median, mean, and trimmed mean) shows comparable results with a Spearman correlation of 0.999 between all methods.

Taxonomic profiles of our urban soil samples were derived from the OTU tables. Differential abundance analysis between cities was performed using MaAsLin3^97^. Features with the false discovery rate (FDR) corrected q-value of the joint prevalence and abundance association <0.1 were considered significant.

## Supporting information

Supplementary Table S1. Metadata and properties of urban soil samples

Supplementary Table S2. Different microbial genera associated with cities and soil properties

Supplementary Table S3. Metadata of urban soil Metagenome-assembled genomes

Supplementary Table S4. Comparison of Species-level genome bins(SGBs)

Supplementary Table S5. The 16S rRNA hits of the MicrobeAtlas database

Supplementary Table S6. Average Nucleotide Identity (ANI) comparison between the longest circular SGB with Rudaea reference genomes

Supplementary Table S7. Biosynthetic gene clusters (BGCs) identified from MAGs

Supplementary Table S8. Small protein families related to the defense system and plasmids

## Supplementary Figures Legends

**Supplementary Fig.1.**
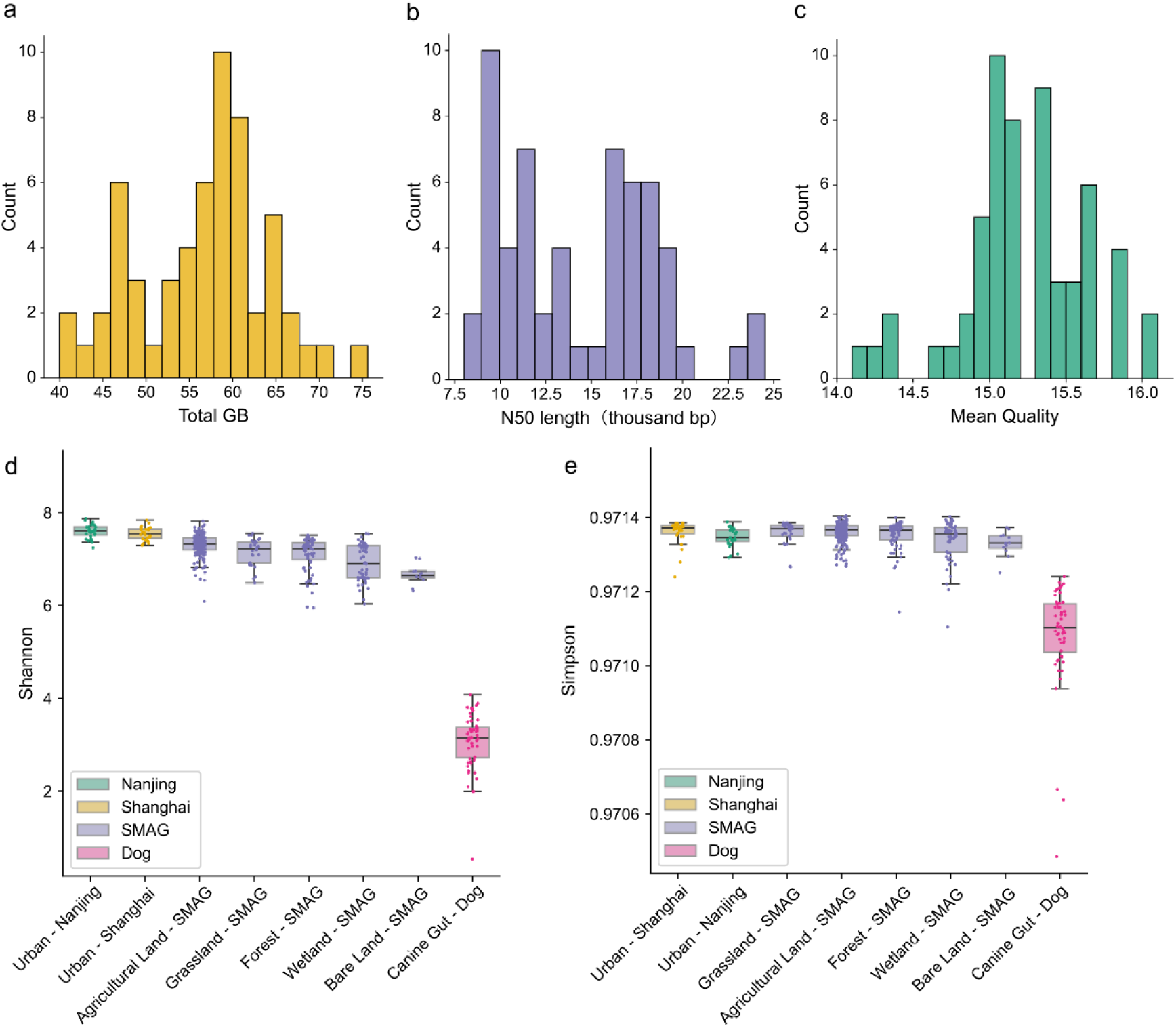
Overview of long reads and microbial alpha-diversity of urban soil samples. **(a)** Sequencing depth of samples. The average is 56.63 Gbp, the maximum is 75.62 Gbp, and the minimum is 39.96 Gbp. **(b)** The N50 length of raw reads from Nanopore sequencing was an average of 14,369 bp, a maximum of 24,526 bp, and a minimum of 8,011 bp. **(c)** The average quality of raw reads from Nanopore sequencing was 15.22, the maximum value was 16.1, and the minimum value was 14.1. **(d-e)** Comparison of the Shannon and Simpson indices between different types of soil, including our samples and those from the SMAG catalog, and a host-associated habitat (pet dog gut). Box plots indicate median (middle line), 25th, 75th percentile (box), and 5th and 95th percentile (whiskers).

**Supplementary Fig.2.**
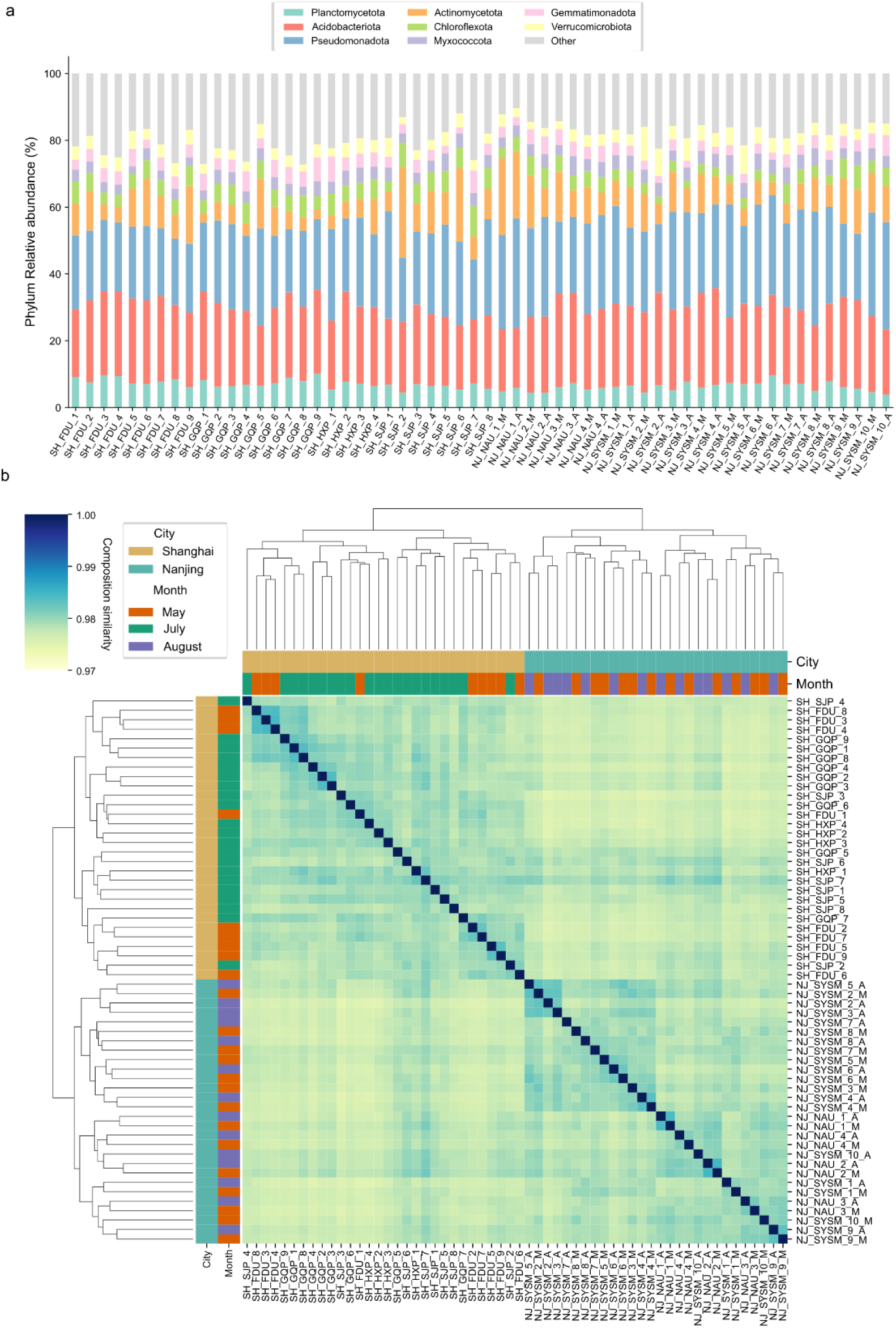
Taxonomic composition and compositional stability of urban soil microbiomes. **(a)** Relative abundance of the top 8 microbial phyla in urban soil samples from Nanjing and Shanghai. **(b)** Compositional similarity of urban soil microbial communities across different sampling sites and time points. SH: Shanghai; NJ: Nanjing; SJP: ShiJi Park; FDU: Fudan University; GQP: GongQing Forest Park; HXP: HuangXing Park; SYSM: Sun Yat-sen Mausoleum; NAU: Nanjing Agricultural University. The sampling months are indicated as M (May), J (July), and A (August).

**Supplementary Fig.3.**
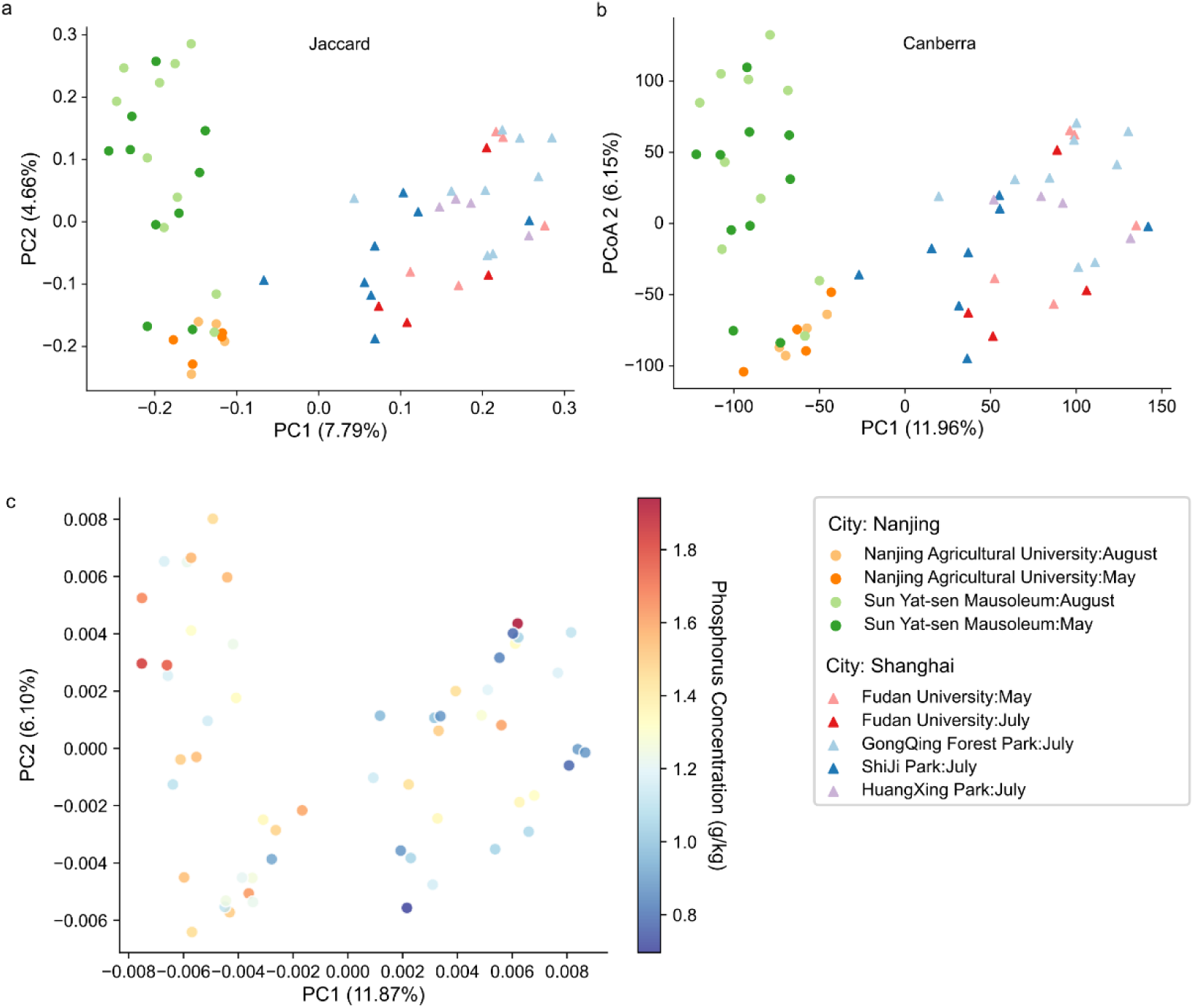
Beta diversity of urban soil microbial communities. **(a-b)** Principal Coordinates Analysis of the beta diversity using the Jaccard distance or Canberra distance. **(c)** Principal Coordinates Analysis of the beta diversity using the Bray-Curtis distance matrix highlights the influence of the soil phosphorus concentration on community clustering, with samples colored according to phosphorus content.

**Supplementary Fig.4.**
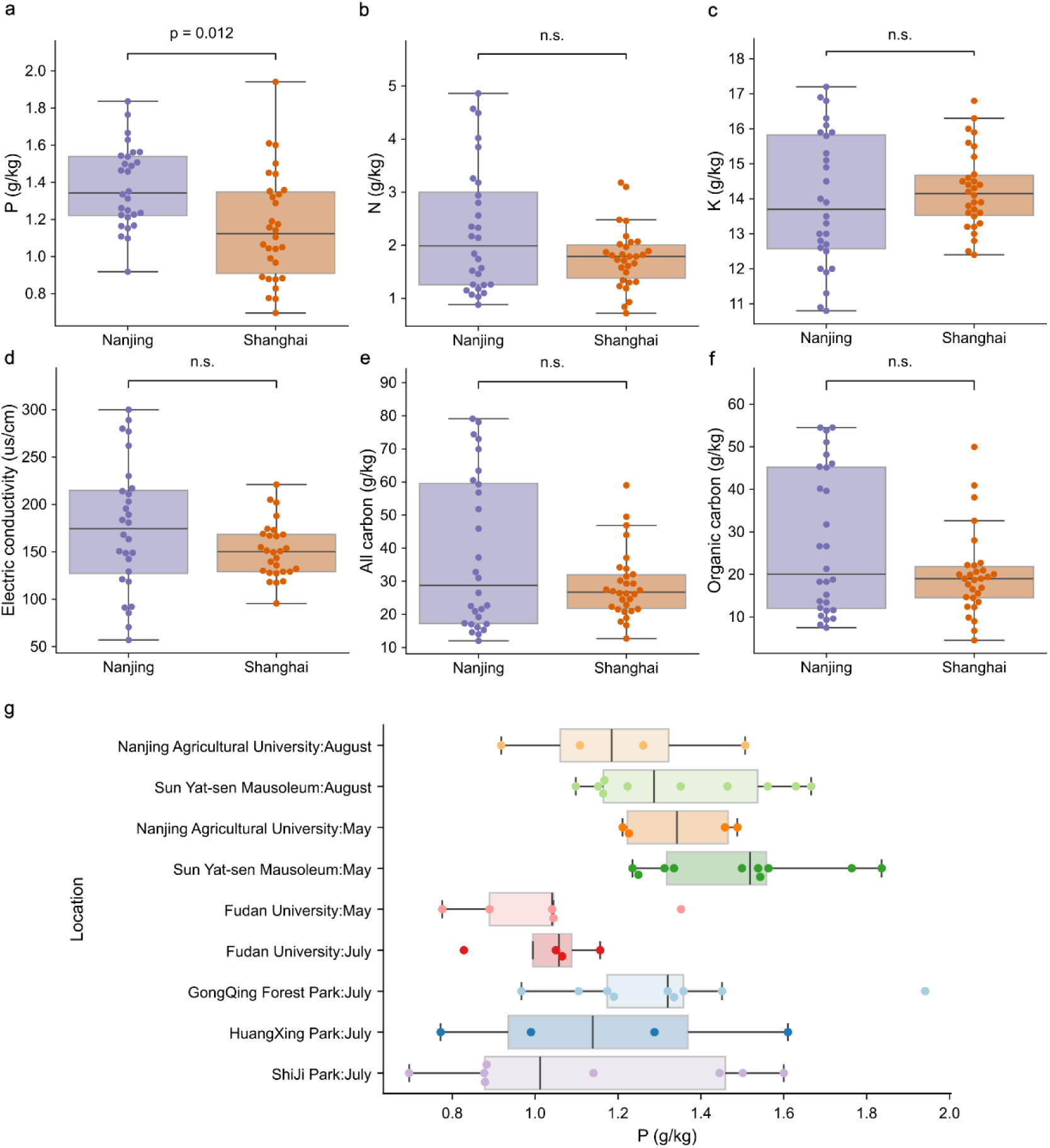
Physicochemical properties at different locations. **(a-f)** Comparison of phosphorus (P), nitrogen (N), and potassium (K) contents; electrical conductivity; total carbon; and organic carbon contents between Nanjing and Shanghai. **(g)** Comparison of phosphorus (P) contents among different locations. Box plots indicate median (middle line), 25th, 75th percentile (box), and 5th and 95th percentile (whiskers).

**Supplementary Fig.5.**
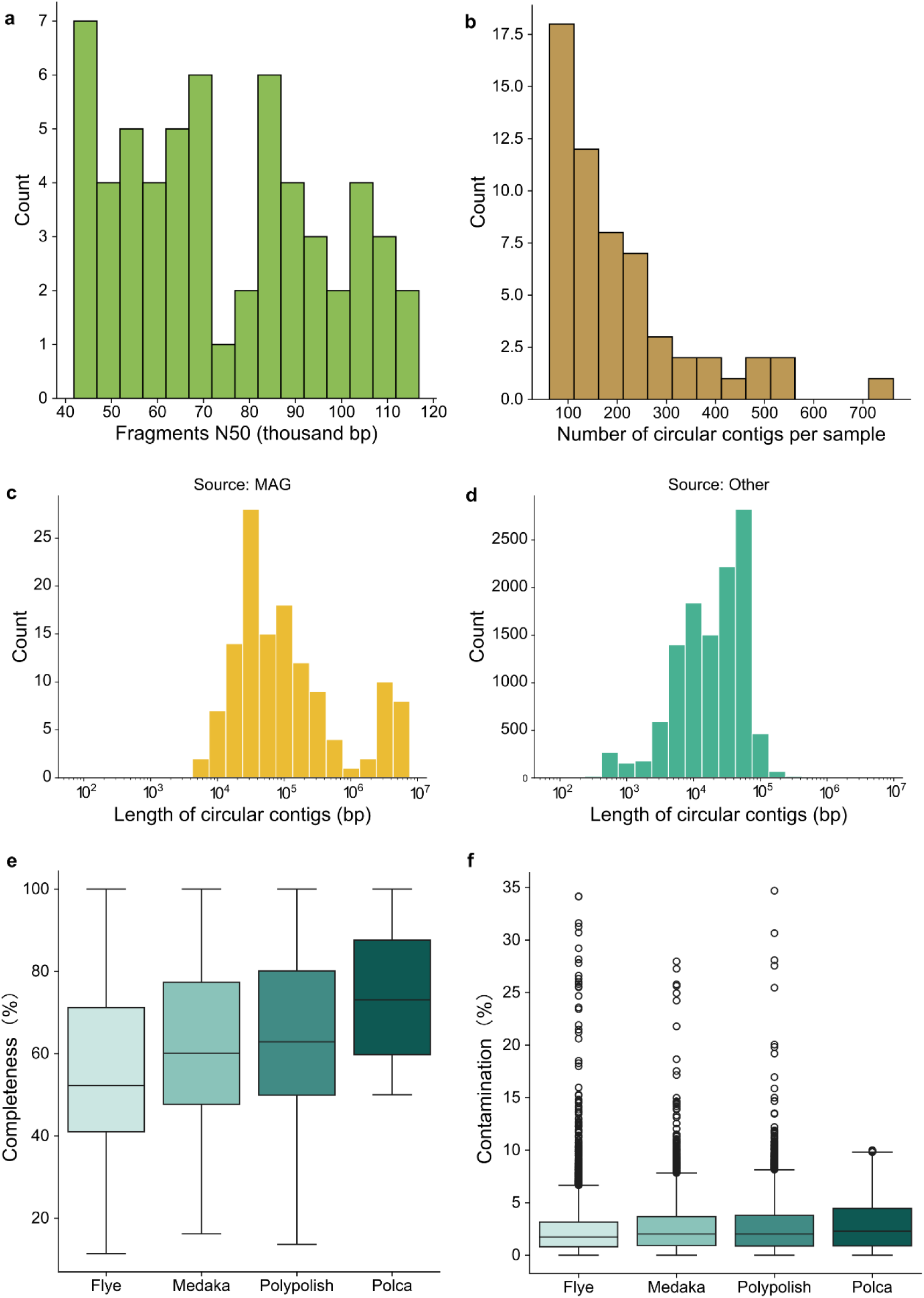
Overview and polishing comparison of assembled contigs. **(a)** The N50 length of the assembled contigs. The average N50 is 74,258 bp, ranging from 41,825 bp to 116,356 bp. **(b)** Distribution of the number of circular contigs assembled per sample. The average number of circular contigs per sample was 201, with a maximum of 750 and a minimum of 62. **(c-d)** Lengths of circular contigs from MAGs or other genetic elements. **(e-f)** Comparison of the completeness and contamination of MAGs after polishing with Medaka, Polypolish, and Polca. Box plots indicate median (middle line), 25th, 75th percentile (box), and 5th and 95th percentile (whiskers) as well as outliers (single points) that lie beyond 1.5 IQRs of the lower and upper quartile.

**Supplementary Fig.6.**
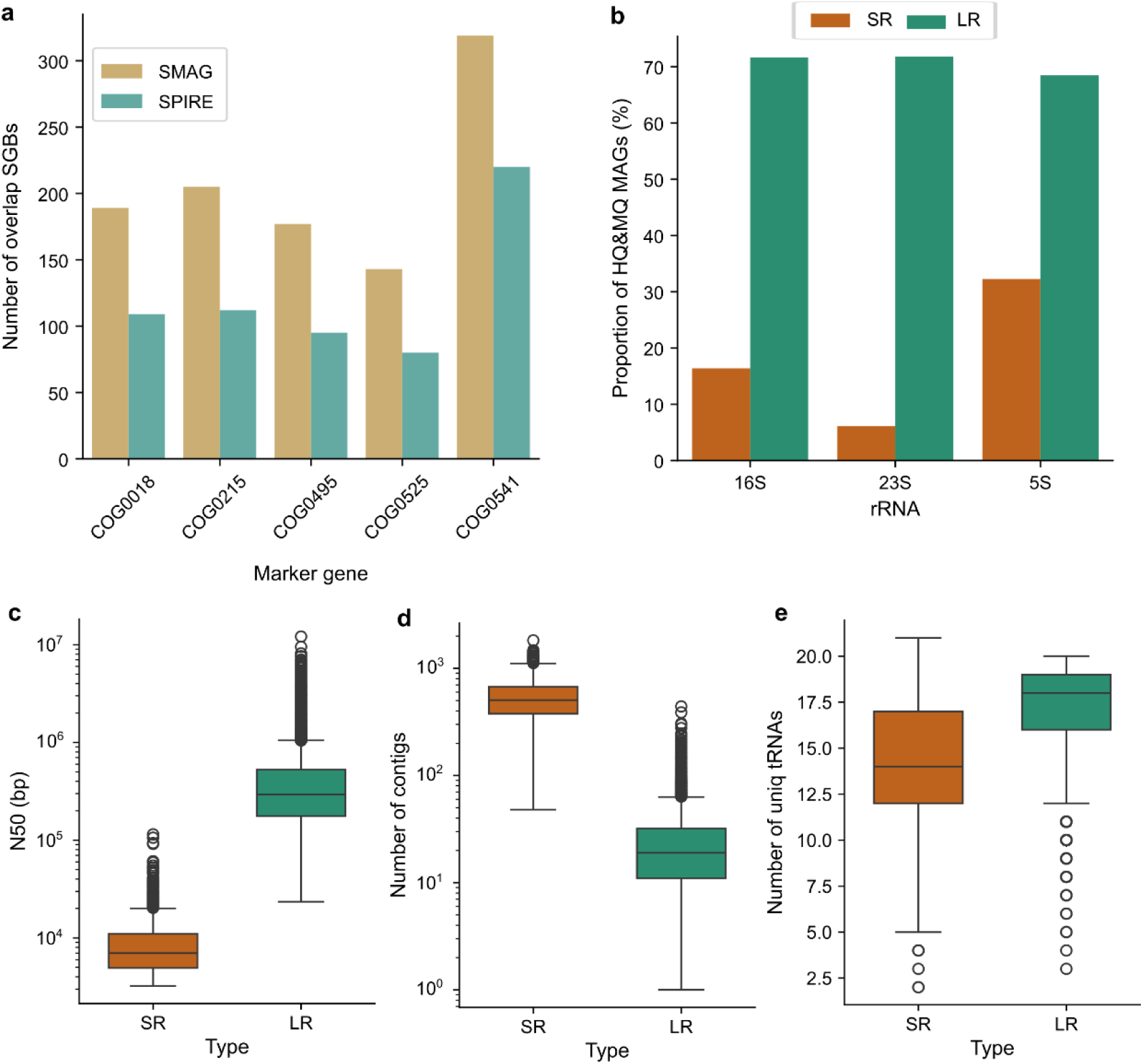
Comparison with other datasets and short-read assemblies. **(a)** Alignment of selected marker genes from 4,171 long-read assembled MAGs and the short-read based SMAG catalog and soil samples in the SPIRE database. **(b)** Comparison of the proportions of long-read assembled and short-read assembled MAGs harboring 16S, 23S, or 5S rRNA. **(c-e)** Comparison of the N50 length, number of contigs, and number of unique tRNAs between long-read assembled MAGs and short-read assembled MAGs. Box plots indicate median (middle line), 25th, 75th percentile (box), and 5th and 95th percentile (whiskers) as well as outliers (single points) that lie beyond 1.5 IQRs of the lower and upper quartile.

**Supplementary Fig.7.**
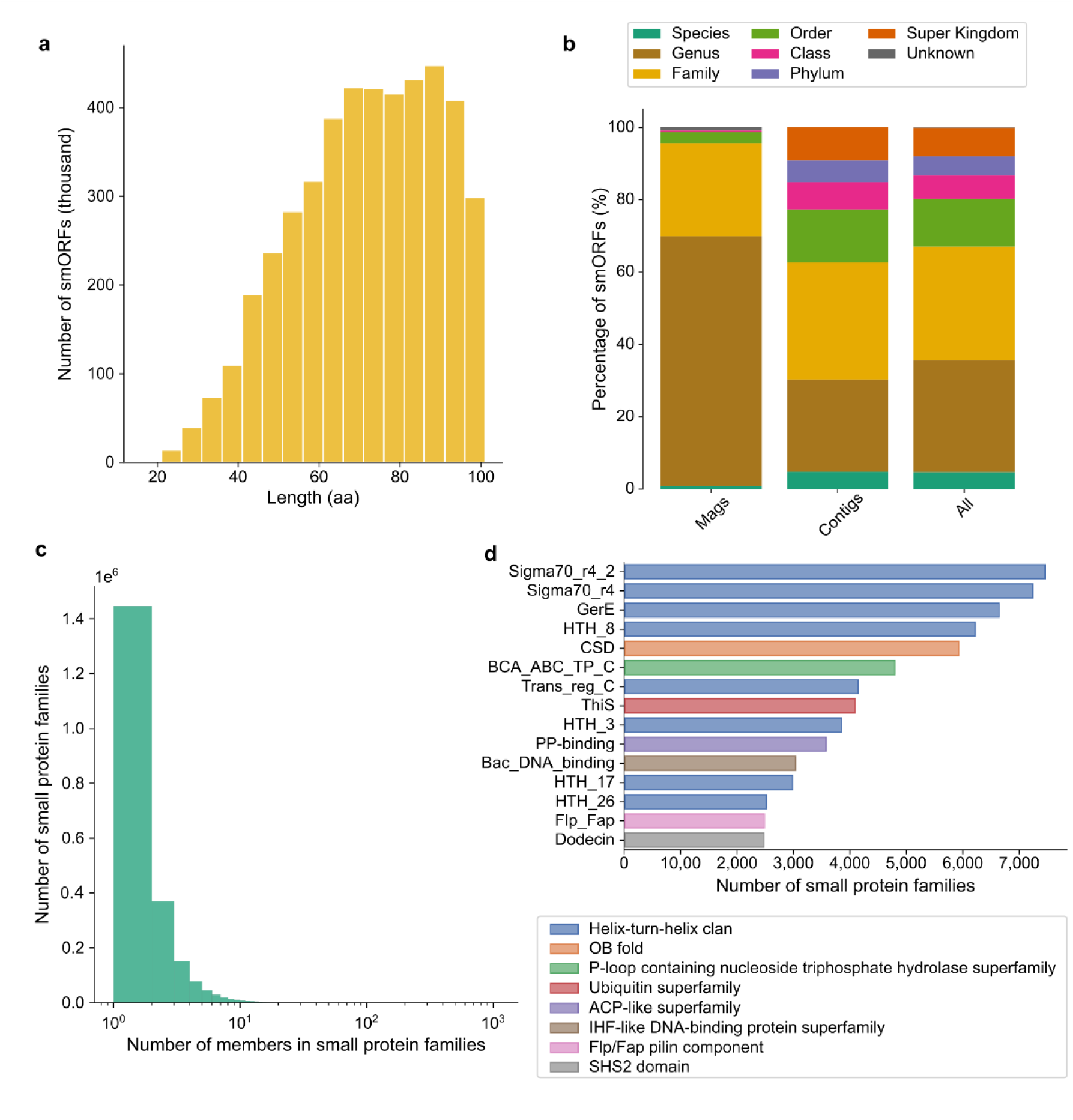
Overview of identified small proteins from the urban soil microbiome. **(a)** The predicted length distribution of small proteins ranges from a minimum of 16 amino acids to a maximum of 99 amino acids. **(b)** Taxonomic classification of predicted smORFs. **(c)** The number of members per small protein family. The largest small protein family contains 1,103 sequences, while 65.5% of the small protein families contain only one sequence. **(d)** The 15 most annotated Pfam conserved domains in the small protein family. The small protein family can be annotated with multiple conserved domains, and multiple counting is used.

**Supplementary Fig. 8.**
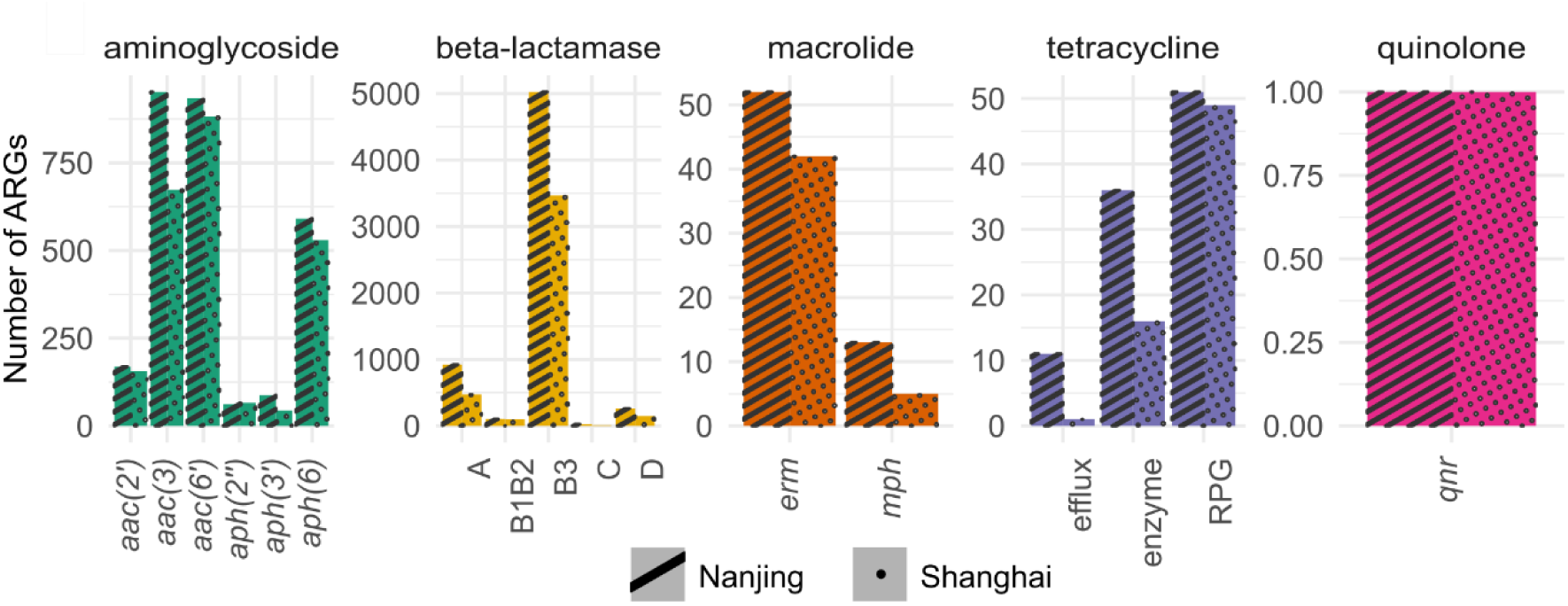
The number of antibiotic resistance genes (ARGs) of urban soil microbiome by gene class. RPG stands for Ribosomal Protection Genes. Striped and dotted columns represent Nanjing and Shanghai, respectively.

## Data Availability

Raw reads and MAG catalog for the urban soil of Shanghai and Nanjing were deposited at the European Nucleotide Archive (ENA) under BioProject ID PRJEB104244.

MAG annotation files have been deposited in Zenodo under DOI: 10.5281/zenodo.18829015

## Code Availability

All original code is available at https://github.com/BigDataBiology/Duan2026_urban_soil.

## Acknowledgements

This work was partly supported by the National Key R&D Program of China (2025YFA1309200, 2023YFF1204800, 2025YFC3409303), the National Natural Science Foundation of China (T2225015, 62433008, 82550127, 42477122, 32250410281), the Shanghai Science and Technology Commission Program (23JS1410100), the Key Science and Technology Project of Hainan Province (ZDYF2024SHFZ058), the Major Project of Guangzhou National Laboratory (GZNL2024A01003), the Fundamental Research Funds for the Central Universities (2025AIYJSZX-YB006), the Natural Science Foundation of Jiangsu Province (BK20230102), the Fundamental and Interdisciplinary Disciplines Breakthrough Plan of the Ministry of Education of China (JYB2025XDXM703), the National Health and Medical Research Council of Australia (under the framework of JPI AMR, #2031902, SEARCHER, to L.P.C.), the Australian Research Council (Future Fellowship, grant #FT230100724, to L.P.C.), and International Development Research Centre, IDRC (under the framework of JPI AMR, grant 109304-001, EMBARK, to L.P.C.). The computations in this research were performed using the CFFF platform of Fudan University.

We thank Alex Chklovski (Queensland University of Technology) for the guidance on plasmid identification and Lisa Galamaga (Queensland University of Technology) for conducting a literature review on soil physical properties that helped inform the interpretation of our results. Members of the Zhao and Coelho group, as well as members of the EMBARK and SEARCHER consortia are thanked for helpful comments during the development of the project.

## Author Contributions Statement

L.P.C. conceptualized and designed the study. Y.D., C.Z., Y.Z., X.Y., and J.Y. collected the samples and the soil-associated information. Y.D., A.Cu., J.I-D., and A.A.C. analysed and visualized the data. L.P.C., X.M.Z., and G.J. supervised the project. Y.D. and L.P.C. wrote the original draft. All authors contributed to the revision of the manuscript prior to submission and approved the final version.

## Competing Interests Statement

A.Cu. is a partner at Nano1Health, SL, and has previously been invited by Oxford Nanopore Technologies (ONT) to participate in conferences. These activities did not influence the results or conclusions of this work. All other authors declare no competing interests.

## Notes

https://doi.org/10.5281/zenodo.18829015

